# High-Quality Synthetic Annotated Tissue Data Using Conditional Generative Adversarial Networks

**DOI:** 10.64898/2025.12.04.690437

**Authors:** Siddharth Srivastava, Cornelis Jan Weijer, Till Bretschneider

**Affiliations:** Department of Computer Science, University of Warwick, Coventry, United Kingdom; School of Life Sciences, University of Dundee, Dundee, United Kingdom

## Abstract

We present a deep-learning based pipeline for generating high-quality synthetic instance-segmentation datasets of tissues undergoing early gastrulation in chick embryo. We create point-clouds using the Lennard-Jones potential and learn an image-to-image translation from these point-clouds to create synthetic tissue images and auxiliary flow tensors using a Generative Adversarial Network, from which the segmentation masks are derived. We evaluate the downstream utility of our synthetic data by training Cellpose and Stardist models from scratch under data replacement and data augmentation scenarios, show that the synthetic datasets effectively capture the statistical properties of the real dataset, and show that our synthetic data improves segmentation performance on a held-out test set. This approach substantially reduces expert annotation time, as late-stage gastrulation data are challenging to acquire and manually label, while similar synthetic examples can be flexibly generated from easily obtainable and annotated early-stage data. Code: https://gitlab.com/siddharthsrivastava/synthetic-tissue-data

## 1 Introduction

The success of deep learning in bioimage analysis has prompted a surge in demand for easily accessible and significant collections of training data to train models to a high degree of accuracy, prompting the exploration of synthetic data generation as a solution for augmenting small, manually annotated training datasets. Traditional data augmentation typically consists of geometric transforms: flips, crops, rotations, translations, scaling, and shearing, to name a few. However, these come with two main problems: firstly, scaling or shearing could change the object’s embedded features, e.g. changing the thickness of a cell membrane in images [1]. Secondly, there is limited control over the augmented training data generated, as we cannot, for instance, increase the diversity of images beyond what is seen in training data.

Several studies have employed Generative Adversarial Networks (GANs) [2] to generate high quality synthetic training data of microscopic cell images [1, 3, 4], predominately to create images of single cells, conditioned on an input shape. Our approach rather introduces a point cloud-conditioned GAN that directly translates the spatial point distributions into full tissue images and their masks, rather than relying on an input noise vector [4, 5] or an entire input shape. This allows explicit control over cell arrangement and density, enabling the generation of statistically realistic epithelial tissue characteristics and alleviating the need for labour-intensive expert annotations of difficult to acquire late-stage data. Our model can be trained on readily obtainable early-stage data to generate realistic late-stage tissues, and preserves biologically meaningful spatial organisation. Additionally, we show that incorporating these GAN-generated images in the training set measurably improves downstream model performance.

## 2 Dataset

We consider images of cells undergoing early-stage gastrulation in the chick embryo, obtained through light-sheet microscopy [6, 7]; we showcase example images in Figure 1.

**Figure 1:**
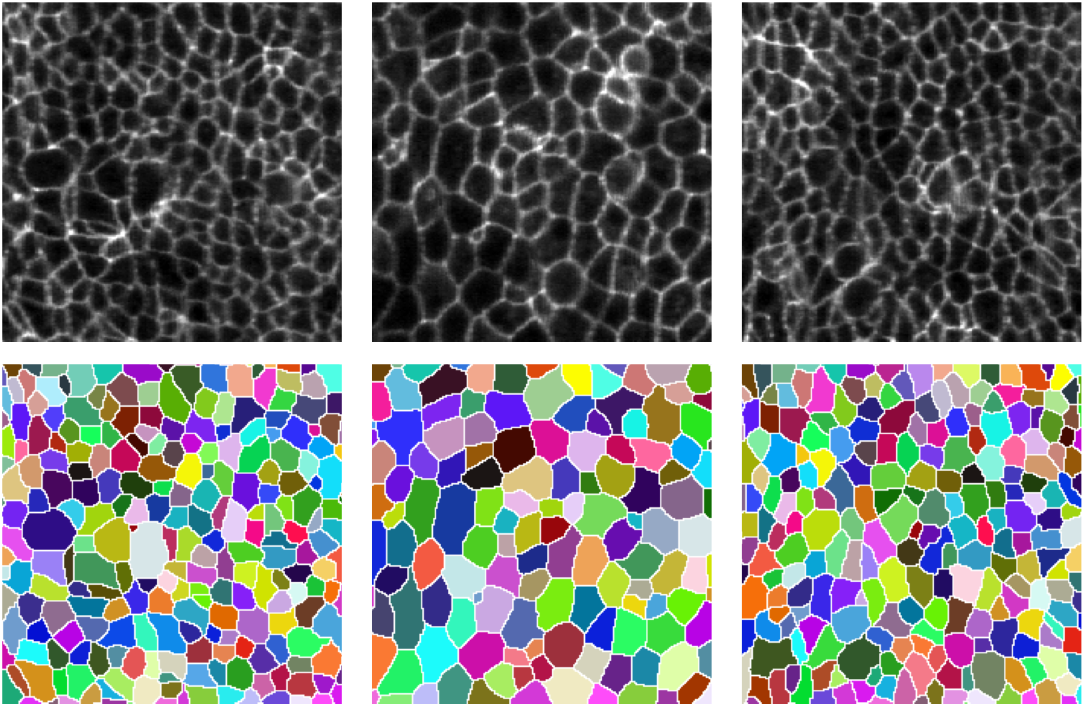
Training set samples: Epithelial tissue images (top) and corresponding ground-truth instance segmentations (bottom, random colour map).

Images are pre-processed using a FFT-based band-pass, a top-hat filter for identifying cell centres, and finally a watershed segmentation is applied to obtain segmented cell boundaries, which is post-processed manually to remove any segmentation artefacts. We focus on early-stage gastrulation as the cells within the tissue are large and easy to segment manually; we showcase that our pipeline provides the flexibility to generate accurate masks for difficult to segment late-stage tissue images. We have labelled images of size 224×224: 98 image-mask pairs for model training and 41 image-mask pairs for testing.

## 3 Methodology

### 3.1 Dataset Preprocessing

We consider a dataset 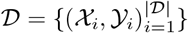 that consists of images 𝒳_*i*_ ∈ ℝ ^*H*×*W*^ and their corresponding segmentation masks 𝒴_*i*_ ∈ ℕ^*H*×*W*^. For each *i* ∈ {1, 2, …, |𝒟|} we define additional auxiliary tensors [*F*_*x,i*_, *F*_*y,i*_, *P*_*i*_, 𝒞_*i*_] as follows:

1. Flow fields: let *D*_*i*_ denote the result of applying a simulated heat diffusion process on 𝒴_*i*_; specifically, for *N* iterations each foreground (i.e. non-zero) pixel is independently assigned the average value of the 3×3 square surrounding it, background pixels are assigned a value of 0, and a constant 1 is added to the centre pixel [8]. We define *F*_*x,i*_ and *F*_*y,i*_ as the discrete horizontal and vertical finite differences of *D*_*i*_, respectively. These are analogous to the horizontal and vertical gradients predicted by Cellpose [8].
2. Binary mask: we define an inside/outside mask *P*_*i*_(*x, y*) such that *P*_*i*_(*x, y*) = 1 if 𝒴_*i*_(*x, y*) *>* 0 and 0 otherwise.
3. Point cloud: let 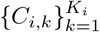 denote the set of connected components of 𝒴_*i*_. For each *C*_*i,k*_, we compute the geometric centre (i.e. centroid) with coordinates 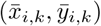. We then define 𝒞 _*i*_ ∈ {0,1}^*H*×*W*^ such that 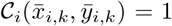, and 0 otherwise.

Therefore 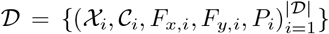 is our final training dataset for the generative model.

### 3.2 Model Architecture

We employ the Wasserstein GAN (WGAN) [9] with gradient penalty (WGAN-GP [10]) as our generative model to create the synthetic dataset. Our generator network consists of two concatenated U-Nets [11] designed to jointly generate the synthetic tissue image and its corresponding segmentation mask.

- The first U-Net extends the unconditional image-to-image translation setup on a noise vector *z* by conditioning on a point cloud 𝒞. A mixture of Gaussian components parameterised by learnable means *µ* and log-variances log *σ* is used to reparametrise *z* [12]:

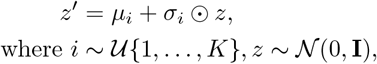

and *K* = 10 is a hyperparameter. The sampled latent vector *z*^′^ is projected through a fully connected layer and concatenated with 𝒞 as input to the U-Net, which outputs three tensors 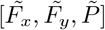. We find that an architecture without *z* results in noticably worse generation.
- The second U-Net takes the generated tensors 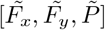 as input and trans-lates them into the final synthetic tissue image 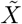 in an image-to-image translation setup.

Therefore, the output of our generator is a 4 × *H* × *W* tensor 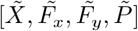of the synthetic flow fields from the first U-Net and the synthetic tissue image from the second U-Net. We term this end-to-end double U-Net setup as a ‘W-Net’. Our discriminator is a Markovian discriminator that outputs a scalar value on the patch level.

### 3.3 Model Training

We train our GAN in the standard Wasserstein training setup; for every generator training step, we train *n*_critic_ = 6 discriminator iterations with the WGANGP loss. We use an additional reconstruction loss of a binary-cross-entropy on the image and *P*-masks and a MSE loss on the flow fields, weighted with *λ* = 10 in the generator loss. We employ the Adam optimiser for both *G* and *D* with *α*_*G*_ = 0.005 and *α*_*D*_ = 0.1 learning rates respectively and instantiate the weights of the generator and discriminator from a *σ* = 0.02 normal distribution. We train for *g*_iter_ = 4000 generator training steps with batch size 16 on a single NVIDIA A40 GPU, taking 4 hours.

### 3.4 Data Postprocessing

To generate the segmentation mask from the GAN outputs, we first compute the divergence field over 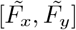 as div 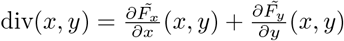. Secondly, we define a set of ‘marker’ regions *M* as the connected components of the sublevel set as *M* := { (*x, y*) : div(*x, y*) ≤1.2}and a foreground mask Ω by the superlevel set of the cell probability map 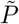 as 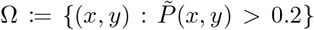, where the 1.2 and 0.2 threshold values were selected through visual inspection. Thirdly, we apply a marker-controlled watershed transform to the divergence field to compute a segmentation *S* := Watershed(div; *M*, Ω) where *M* is used as the object basins for the watershed, and Ω is used to constrain the watershed to only label foreground cell pixels. Finally, the connected components in the output segmentation *S* are assigned unique integer labels using a connected components labelling algorithm to compute the final 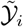. The generated tissue image 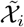 is replaced by a histogram-matched version of itself with respect to 𝒳_*r*(*i*)_, where *r*(*i*) ~ 𝒰 {1, …, |𝒟|}, i.e. a randomly sampled real image. This is such that the intensity distribution of our synthetic images match that of the real images. We visually showcase the stages of this post-processing in Figure 2.

**Figure 2:**
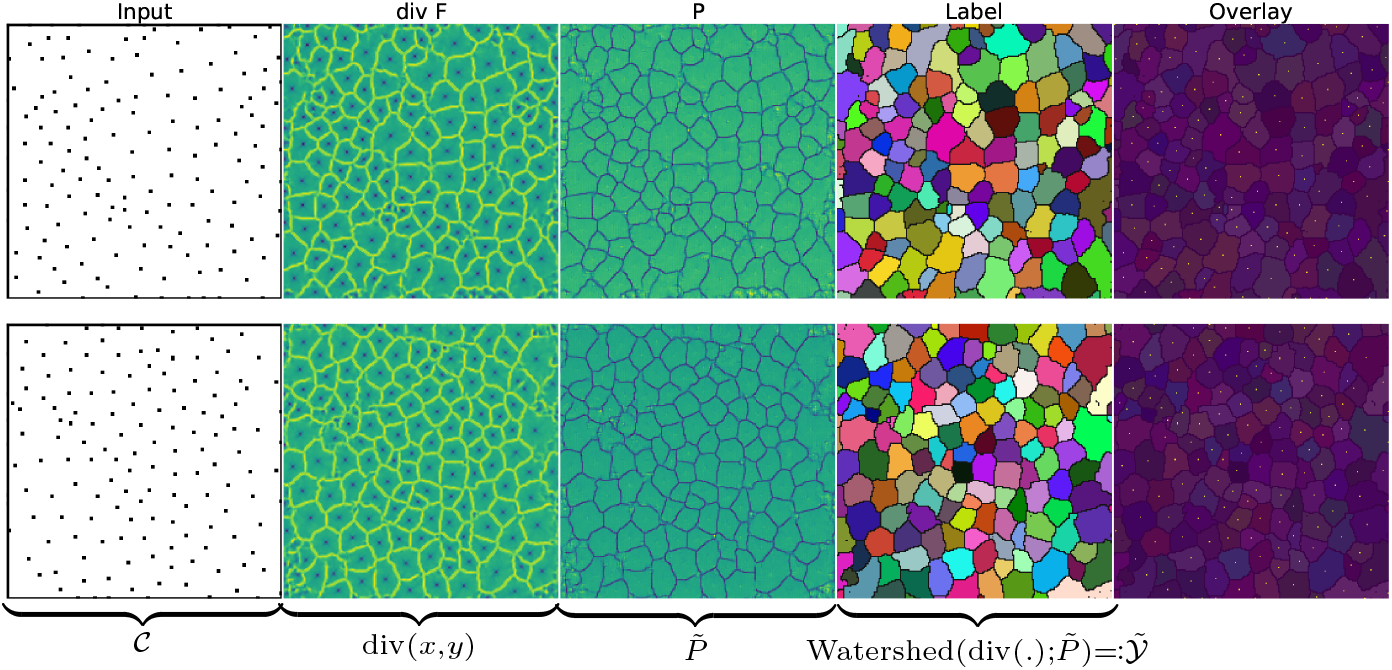
Postprocessing *G*(𝒞) to generate the label 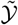 corresponding to 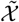. The overlay of 𝒞 over the label mask confirms the GAN learns to map each point in 𝒞 to the centroid of a cell within the generated tissue. Size of points in the 𝒞 has been increased for visualisation.

### 3.5 Point Cloud Generation

The GAN architecture described earlier translates an input binary point-cloud 𝒞 _*i*_ ∈ {0,1}^*H*×*W*^ into our intermediary tensors, which are post-processed to generate the final synthetic dataset. To generate these point clouds, we use a model based on the Lennard–Jones 12-6 (LJ) potential [13], which defines a pairwise interaction potential between particles *i* and *j* as

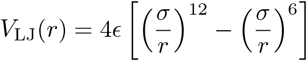

where *r* is the Euclidean distance between particles *i* and *j*, and *ϵ, σ* are hyper-parameters. The number of points *n* is (independently for each mask) sampled from a kernel density estimate of the empirical distribution of particle counts in the training data; the *n* particles’ coordinates are initialised randomly and optimised via gradient-based minimisation of the sum over all *V*_LJ_(*r*) for *K* = 1000 iterations. We choose the learning rate of this minimisation as *α* = 0.005 and set *σ* = 0.2, *ϵ* = 0.5 in *V*_LJ_.

## 4 Experiments and Results

### 4.1 Visual Accuracy & Shape Characteristics

We provide representative examples from our synthetic dataset in Figure 4, showcasing synthetic tissues generated from sparse point clouds in Figure 4a and tissues from dense point clouds in Figure 4b. Our method generates visually realistic tissue images when compared to the real images in Figure 1 with minimal artefacts in both the tissue image and the segmentation mask. Visual inspection confirms strong correspondence between each image and its mask, indicating that the generated annotations are well aligned with the corresponding tissue structure. We investigate further by comparing the shape characteristics of the cells within the synthetic and real tissue images in Figure 3, looking at the distributions of eccentricity, solidity, orientation, and perimeter of all cells in the dataset. We show that our GAN successfully captures the underlying tissue shape characteristics in the real dataset, through the strong alignment between the real and synthetic histograms in terms of overlapping distributions and similar peak structures, and demonstrating that the GAN is not generating arbitrary or uniform shapes, but rather captures the statistical and morphological properties of the real tissue.

**Figure 3:**
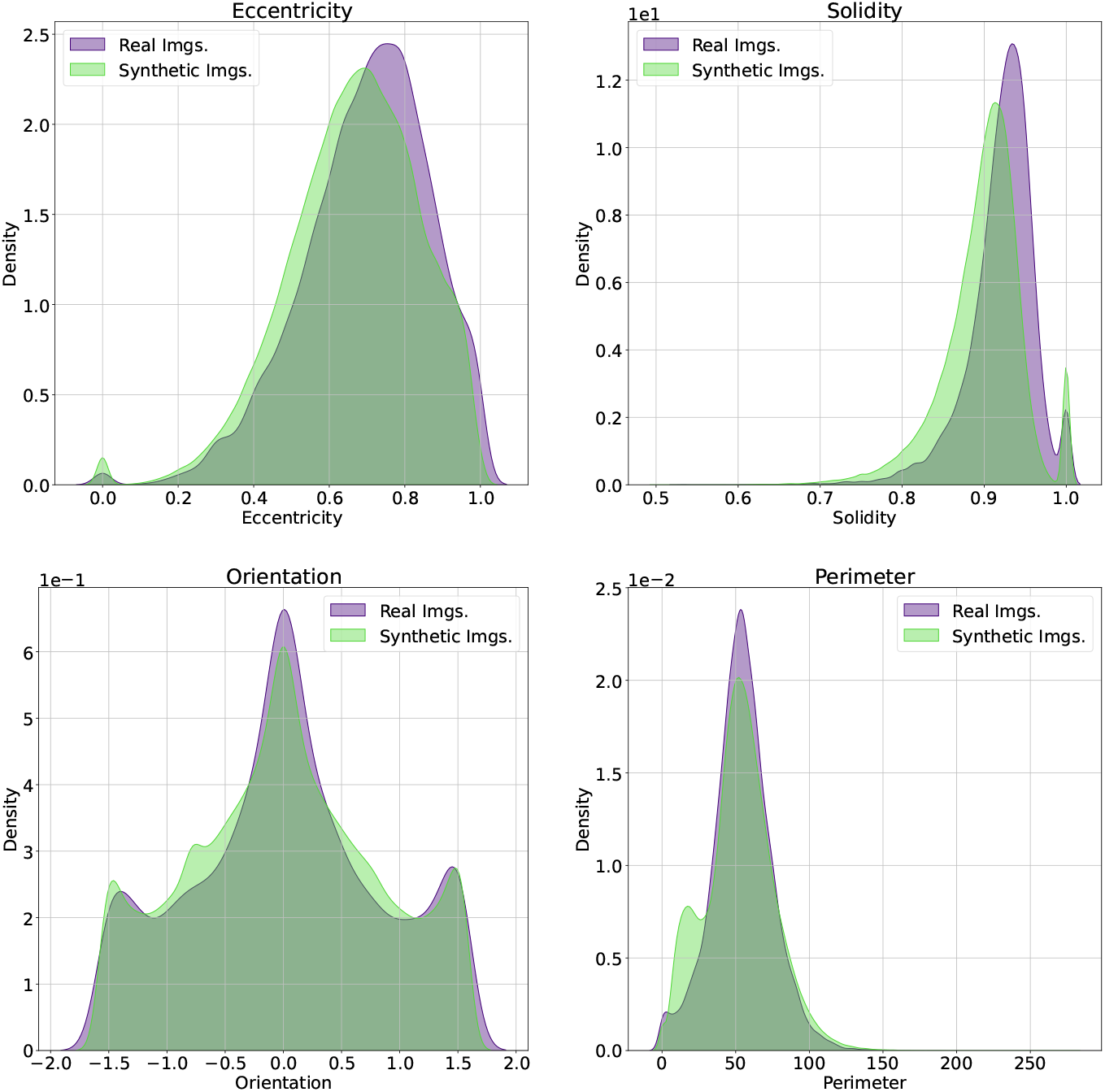
KDE plot of the eccentricity, solidity, orientation and perimeter shape descriptors between real (purple) and GAN-generated (green) tissue instance segmentation masks. Synthetic densities are computed using 10 *×* | 𝒟| tissue images.

**Figure 4:**
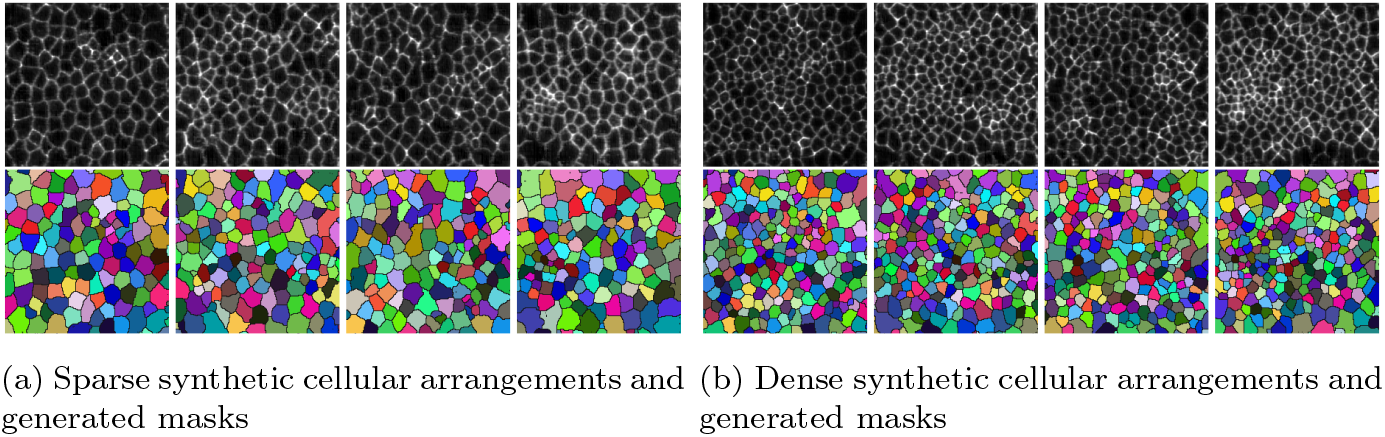
Sparse (left) and dense (right) synthetic tissue arrangements and their corresponding instance segmentation masks.

### 4.2 Downstream Experimental Setup

We train two popular models for cell segmentation, Cellpose [8] and Stardist [14], from scratch using varying proportions of real and synthetic data, and evaluate performance on a held-out real test set. Each model was trained for 300 epochs across ten independent runs with default settings. We consider two experimental settings:

- **Data Replacement**. The original training dataset is wholly replaced with varying proportions of synthetic data - specifically, 1×, 3×, and 7× the number of real samples. Model performance is subsequently assessed on a held-out real test set. This quantifies the utility of the generated samples to serve as an effective replacement for the real data in model training.
- **Data Augmentation**. The synthetic data are used to expand the real training set by augmenting it with additional synthetic samples corresponding to 1×, 3×, and 7 × the size of the real dataset. This is a more realistic situation of quantifying whether synthetic data together with realdata can enhance generalisation and robustness.

We showcase our experimental results in Figure 5. In the ‘Replace’ experiments, both models maintain comparable Jaccard scores to those trained only on the real dataset, indicating the effectiveness of our synthetic dataset in capturing the statistical properties of underlying biological structures from the real dataset. Data augmentation proves measurably beneficial for Cellpose, increasing segmentation performance with only 1× synthetic data, suggesting that our synthetic data improves model generalisation. Synthetic data proved less beneficial for Stardist. These results demonstrate that our data generation approach not only provides a viable substitute for manually annotated data but also contributes to stronger model performance through dataset augmentation.

**Figure 5:**
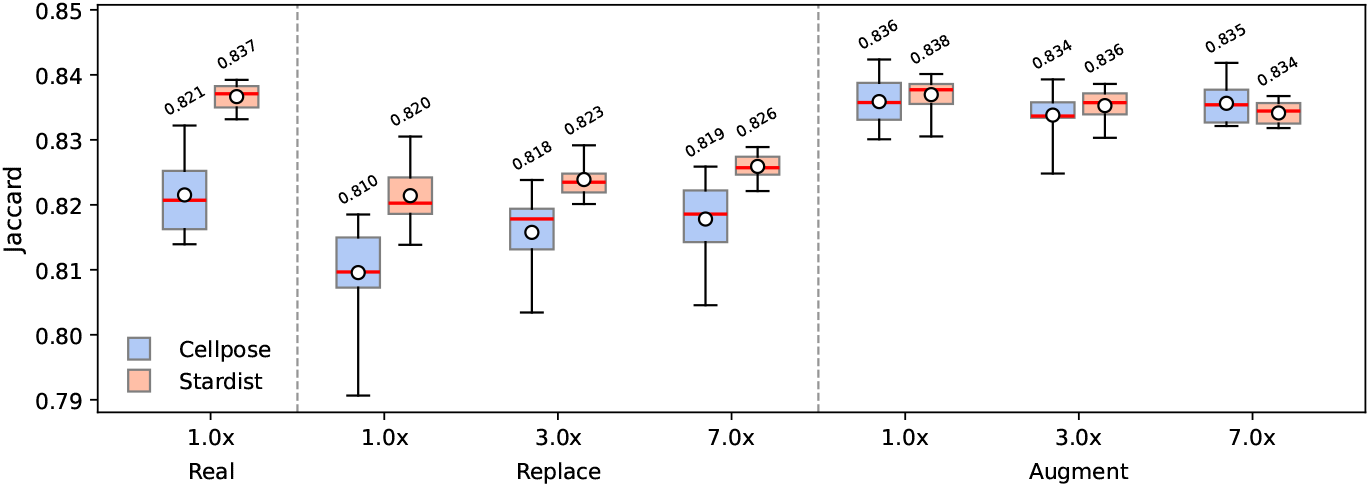
Box plots showing Cellpose and Stardist performance under Replacement and Augmentation scenarios with varying amounts of real and synthetic data, evaluated on a held-out real test set (10 independent trials). Boxes show the 25–75% range; whiskers the min–max; red line the median; white circle the mean. Median values are shown above each box.

## 5 Concluding Remarks

This paper presents a pipeline for generating synthetic tissue images and segmentation masks using deep generative models, and showcases that the synthetic dataset captures the key statistical properties of real microscopy data. We confirm downstream utility of our data by training segmentation models on the synthetic dataset and testing on the real dataset, demonstrating strong and improved performance. A major contribution is our point cloud to segmentation translation and showcasing the robustness of the pipeline across different cell densities within the tissue. Using point clouds as input gives fine-grain control over the tissue architecture that we wish to generate and allows balancing a dataset, for example to enrich it with early dividing cells, which are larger and more round. Owing to its flexibility, our approach is broadly applicable to diverse tissue imaging modalities, developmental stages, and naturally lends itself to transfer learning as many tissue architectures are highly similar. This will reduce the amount of manually labelled training data which will only be needed to inform specific fluorescence distributions, offering a scalable alternative to manual annotation and data acquisition.

## 6 Compliance with ethical standards

This is a numerical simulation study for which no ethical approval was required.

## 7 Acknowledgments

The authors acknowledge the use of the Batch Compute System in the Department of Computer Science at the University of Warwick, and associated support services, in the completion of this work. T.B. and S.S. acknowledge support through EPSRC grants EP/V062522/1 and EP/X026663/1.

## References

[1] Piotr Baniukiewicz, E. Josiah Lutton, Sharon Collier, and Till Bretschneider, “Generative Adversarial Networks for Augmenting Training Data of Microscopic Cell Images,” in Frontiers in Computer Science, 2019, vol. 1.

[2] Ian J. Goodfellow, Jean Pouget-Abadie, Mehdi Mirza, Bing Xu, David Warde-Farley, Sherjil Ozair, Aaron Courville, and Yoshua Bengio, “Generative Adversarial Nets,” in Proceedings of the 28th International Conference on Neural Information Processing Systems, 2014.

[3] Peter Goldsborough, Nick Pawlowski, Juan C Caicedo, Shantanu Singh, and Anne E Carpenter, “Cytogan: Generative modeling of cell images,” bioRxiv, 2017.

[4] Kevin Barrera, Anna Merino, Angel Molina, and José Rodellar “Automatic generation of artificial images of leukocytes and leukemic cells using generative adversarial networks (syntheticcellgan),” Comput. Methods Prog. Biomed., 2023.

[5] John T. Guibas, Tejpal S. Virdi, and Peter S. Li, “Synthetic Medical Images from Dual Generative Adversarial Networks,” in Proceedings of the 31st Conference on Neural Information Processing Systems (NIPS 2017), 2017, vol. NeurIPS ‘17.

[6] Emil Rozbicki, Manli Chuai, Antti Karjalainen, Feifei Song, Helen M. Sang, Rene Martin, Hans-Joachim Knolker, Michael MacDonald, and Cornelis J. Weijer, “Myosin-II-mediated cell shape changes and cell intercalation contribute to primitive streak formation,” Nature Cell Biology, 2015.

[7] Manli Chuai, Guillermo Serrano Nájera, Mattia Serra, Lakshminarayanan Mahadevan, and Cornelis J. Weijer, “Reconstruction of distinct vertebrate gastrulation modes via modulation of key cell behaviors in the chick embryo,” Science Advances, vol. 9, no. 1, 2023.

[8] Carsen Stringer, Tim Wang, Michalis Michaelos, and Marius Pachitariu, “Cellpose: A generalist algorithm for cellular segmentation,” Nature Methods, vol. 18, no. 1, Jan. 2021.

[9] Martin Arjovsky, Soumith Chintala, and Léon Bottou, “Wasserstein generative adversarial networks,” in Proceedings of the 34th International Conference on Machine Learning - Volume 70, 2017, ICML’17.

[10] Ishaan Gulrajani, Faruk Ahmed, Martin Arjovsky, Vincent Dumoulin, and Aaron Courville, “Improved training of wasserstein GANs,” in Proceedings of the 31st International Conference on Neural Information Processing Systems, 2017.

[11] Olaf Ronneberger, Philipp Fischer, and Thomas Brox, “U-Net: Convolutional Networks for Biomedical Image Segmentation,” in Medical Image Computing and Computer-Assisted Intervention – MICCAI 2015, 2015.

[12] Swaminathan Gurumurthy, Ravi Kiran Sarvadevabhatla, and R. Venkatesh Babu, “DeLiGAN: Generative Adversarial Networks for Diverse and Limited Data,” in 2017 IEEE Conference on Computer Vision and Pattern Recognition (CVPR), 2017.

[13] Xipeng Wang, Simón Ramírez-Hinestrosa, Jure Dobnikar, and Daan Frenkel, “The Lennard-Jones potential: When (not) to use it,” Physical Chemistry Chemical Physics, 2020.

[14] Alejandro F. Frangi, Julia A. Schnabel, Christos Davatzikos, Carlos Alberola-López, Gabor Fichtinger, Uwe Schmidt, Martin Weigert, Coleman Broaddus, and Gene Myers, “Cell Detection with Star-Convex Polygons,” in Medical Image Computing and Computer Assisted Intervention – MICCAI 2018, 2018.

